# Integrating independent microbial studies to build predictive models of anaerobic digestion inhibition by ammonia and phenol

**DOI:** 10.1101/2020.03.16.993220

**Authors:** Simon Poirier, Sébastien Déjean, Cédric Midoux, Kim-Anh Lê Cao, Olivier Chapleur

## Abstract

Anaerobic digestion (AD) is a microbial process that can efficiently degrade organic waste into renewable energies such as methane-rich biogas. However, the underpinning microbial mechanisms are highly vulnerable to a wide range of inhibitory compounds, leading to process failure and economic losses. High-throughput sequencing technologies enable the identification of microbial indicators of digesters inhibition and can provide new insights into the key phylotypes at stake during AD process. But yet, current studies have used different inocula, substrates, geographical sites and types of reactors, resulting in indicators that are not robust or reproducible across independent studies. In addition, such studies focus on the identification of a single microbial indicator that is not reflective of the complexity of AD. Our study proposes the first analysis of its kind that seeks for a robust signature of microbial indicators of phenol and ammonia inhibitions, whilst leveraging on 4 independent in-house and external AD microbial studies. We applied a recent multivariate integrative method on two-in-house studies to identify such signature, then predicted the inhibitory status of samples from two datasets with more than 90% accuracy. Our study demonstrates how we can efficiently analyze existing studies to extract robust microbial community patterns, predict AD inhibition, and deepen our understanding of AD towards better AD microbial management.

**Highlights:**

- Robust biomarkers of AD inhibition were tagged by integrating independent 16S studies
- Increase of the *Clostridiales* relative abundance is an early warning of AD inhibition
- *Cloacimonetes* is associated with good performance of biomethane production
- Multivariate model predicts ammonia inhibition with 90% accuracy in external data

## 1 Introduction

Anaerobic digestion (AD) is considered as the most efficient and sustainable technology for organic waste treatment. It has the ability to enable simultaneously waste stabilization and valorization through the production of methane rich biogas and of digestate used as an organic amendment. Encouraged by the renewable energy policies, biogas production with anaerobic digestion has increased in the EU to reach 18 billion m^3^ methane (654 PJ) in 2015 [1]. However, The European Biomass Association (AEBIOM) estimates that anaerobic digestion still has a considerable potential for expansion with a biogas potential at about 78 billion m^3^ biomethane. To reach this goal, the optimization of biogas production is essential to improve high process stability and efficiency and lower susceptibility to disturbances. Indeed, process failure reduces the economic and environmental performances of biogas technology as they lead to decreased methane yields and thus reduce revenues. Therefore, it is important that applied research on biogas technology improve robustness of these systems to stress factors, such as altered operating conditions or inhibitory compounds.

Among the broad range of inhibitors from AD substrates, high concentrations of ammonia and micro-pollutants such as phenol are considered as the primary cause of digester failure [2, 3]. Commonly used feedstock such as livestock manure, slaughterhouse byproducts and food industrial residues contain organic nitrogen such as urea and proteins, which readily release ammonia during their anaerobic degradation [4]. In addition, various natural or anthropogenic phenolic compounds are detected in different types of effluents from coal gasification, coking, petroleum refining, petrochemical manufacturing and paper [5]. Phenols are also produced from biodegradation of naturally occurring aromatic polymers such as humic acids and tannins or from degradation of xenobiotic compounds such as pesticides [6]. As a result, contaminated sludge produced during the treatment of these various effluents can cause digester imbalance.

The stability and efficiency of the overall AD process relies on tightly coupled synergistic activities between an intricate community of microorganisms. But the understanding of biological mechanisms of AD is still hampered by the extreme complexity of the microbial ecosystem involved in this process [7, 8]. New knowledge is needed to unravel bioindicators of digesters inhibition which have the potential to guide and optimize operation management during unexpected onset of inhibitors and prevent biogas production.

High-throughput sequencing technologies, including 16S rRNA amplicon sequencing have enabled many studies to monitor microbial changes during steady state [9-11] or the inhibition of anaerobic digesters, for example with phenolic compounds [12-14] or ammonia [15-18]. However, given a same inhibitor, independent studies identified very different microbial indicators, due to differences in inocula and substrates usage, geographical sites and/or at different times, and types of digesters. The lack of reproducibility is further accentuated when studies focus on the identification of a single microbial bioindicator. Such univariate perspective is unlikely to shed light into the global and complex ecosystem of AD.

Our study aims at identifying a robust multivariate microbial signature reflective of the interaction network within anaerobic digesters whilst leveraging on in-house and other existing studies. The analytical challenge was to combine such independent studies plagued by unwanted variation (e.g. different substrates and types of digesters) that outweigh the interesting biological variation of ammonia and phenol inhibitions. We applied a recently developed integration method MINT (Multivariate INTegrative) [19] that provides an integrated view of anaerobic digester microbiota subject to distinct types of inhibition. By integrating two independent in-house experiments, we identified robust microbial bioindicators that characterize ammonia inhibition, phenol inhibition and no inhibition. We evaluated this model by predicting AD status on two external studies assessing the influence of ammonia on AD. By doing so, we demonstrate the feasibility of detecting robust indicators evidencing the microbial symptoms of AD process dysfunction in independent studies which suggests promising applications in various biotechnologies thanks to the expansion of data deposited in public databases.

## 2 Materials and methods

### 2.1 Experimental data

Four studies assessing the influence of ammonia or phenol on anaerobic digestion were selected to build the predictive models [20-23]. In all the studies, gas productions were measured to evaluate if the anaerobic digestion was inhibited. As described in Table 1, two out of four studies were conducted in our laboratory with the same type of substrate (biowaste) but with two different inocula collected one year apart from an industrial mesophilic digester treating wastewater treatment sludge. Samples were taken across time and under different inhibitory conditions. DNA was extracted and 16S rRNA gene was sequenced providing datasets of raw sequences associated to different inhibitory conditions. Study 1 aimed at assessing in parallel the effect of different levels of total ammonia nitrogen (TAN) and phenol on the microbial community of batch anaerobic digester [21]. Study 2 assessed the influence of support media addition to mitigate anaerobic digestion ecosystem inhibition in presence of two inhibitory conditions (19 g/L of TAN and 1.5 g/L of phenol, respectively) [20]. Studies 3 and 4 were conducted in two distinct external laboratories. Study 3 conducted by Lü *et al.* evaluated the effectiveness of biochar of different particle sizes in alleviating different ammonia inhibition levels during anaerobic digestion of 6 g/L glucose [23]. Study 4 by Peng et al. sought for microbial community changes during inhibition by ammonia in high solid anaerobic digestion of food waste in a continuous stirred-*tank* reactor [22].

**Table 1:**
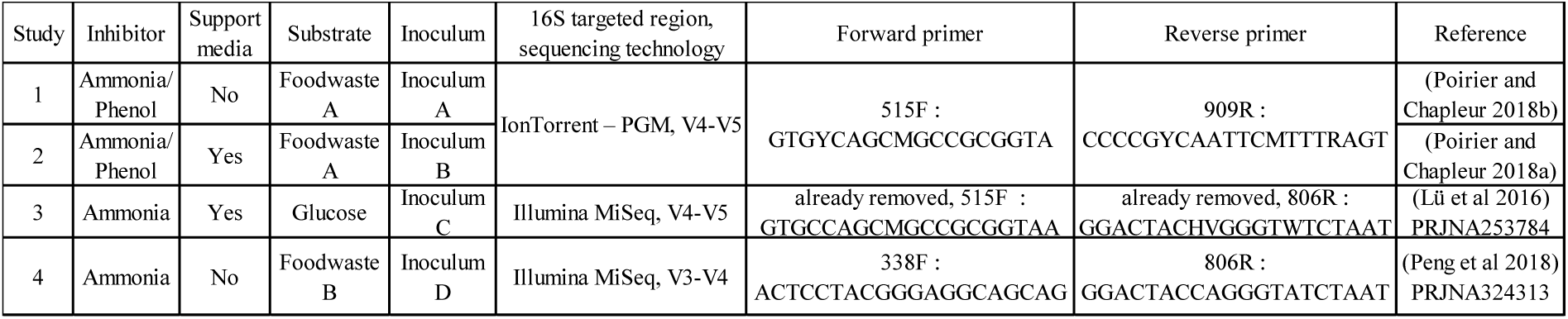
Description of the datasets used for the analysis

In order to train an accurate and relevant our MINT model before prediction, we removed some samples from studies 1 and 2 that were deemed non-representative of our analytical objectives. They are listed in supplementary material (table S1). In studies 1 and 2, only samples collected after at least 10 days of incubation were kept to ensure that the microbial community was representative of the inhibitory conditions. Samples taken after more than 60 days of incubation were removed as biogas production was completed. Moreover, for studies 1 and 2, methane cumulated production data were fitted to a modified Gompertz three-parameter model (Eq. (1)) where M(t) is the cumulative CH_4_ production (mL) at time t (d); P is the ultimate CH_4_ yield (mL); R_max_ is the maximum CH_4_ production rate (mL/d); λ is the lag phase (d); e is the exponential:

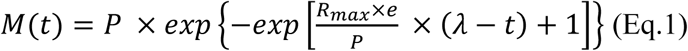

Reactors were deemed inhibited when Rmax was less than 80% of Rmax in the controls without inhibitor and not inhibited when Rmax was at least 90% of Rmax in the controls without inhibitor. In study 1, samples from reactors incubated with 250 or 500 mg/L of phenol and 5.0 g/L of ammonium were discarded as the inhibition status (inhibited-non-inhibited) was not well defined. Samples collected from batch digesters incubated with 50 g/L of ammonium or 5 g/L of phenol were removed as these incubations were totally inhibited and microbial community did not evolve. In total 81 samples remained in studies 1 and 2. All samples from study 3 and bacterial samples of study 4 were kept to be tested with the model (respectively 37 and 10, table S1).

### 2.2 Data processing

In these four studies, sequencing of the V3-V4 or V4-V5 region of the 16S rRNA gene had been performed with three different approaches, as described in Table 1. Data from external studies were downloaded from NCBI with fastq-dump 2.8.1. Paired-end reads from Lü *et al*., and Peng *et al.*, studies were merged with pear v0.9.11 [24]. Adapters from each study were specifically removed with cutadapt v1.12 [25]. All sequences were imported into FROGS pipeline [26]. Samples from studies 1 and 2 were processed together while studies 3 and 4 were processed independently because of the differences in sequencing approaches. Taxonomic assignment of OTUs was performed using Silva 132 SSU as reference database. OTUs were trimmed by keeping only those present more than 10 times in the whole dataset (resp. 1133, 399, 158 OTUs for studies 1 and 2, 3, 4). For joint analysis of data from studies 1 and 2, data was processed as obtained. For joint analysis of studies 1, 2, 3 and 4, the three distinct biom files were concatenated and data were discussed at the genus level. Sequences of interest were then assigned at the species level using the Blastn+ algorithm [27].

### 2.3 Statistical analyses and predictive model

OTUs abundances were scaled with total sum scaling to account for uneven sequencing depth. OTUs that exceeded 3% in at least one sample were retained for the analysis. The total relative abundance of these minor OTUs represented 17% of the total number of sequences. Data were then transformed with centered log ratio (CLR) transformation to account for compositional structure of the scaled data. All statistical analyses were implemented with mixOmics R package, as described in [28].

In order to obtain a first understanding of the major sources of variation in the training data (studies 1 and 2), and to obtain a first insight into the similarities between samples, we conducted principal component analyses (PCA) on the 16S rRNA tags datasets (pca() function). A sparse Partial Least Squares Discriminant Analysis (Sparse PLS-DA) was then conducted to assess the potential to discriminate the samples according to the type of inhibition [29] (sPLS-DA() function) and identify microbial signatures characterizing inhibition type. Classification accuracy was calculated based on the microbial signature identified by the method, as described in [28]. Finally, the MINT sPLS-DA method (referred to as MINT in the following), that generalizes sPLS-DA while accounting for study-specific effects was applied [19] (mint.splsda() function). Parameters to choose in MINT included the number of PLS-DA components, and the number of variables to select, which was performed using 10-fold cross-validation. The final MINT model was then fitted on the data, and the classification performance was estimated using the perf() function and 10-fold cross-validation repeated 10 times. Graphical display of the discriminative OTU signature identified by MINT were output using clustered image maps (cim() function). The multivariate model not only identifies a microbial signature characterizing inhibition status, but it also enables to predict the groups of samples from external data sets as described in detail in [28]. For prediction, we trained a new MINT model on studies 1 and 2 for conditions ammonia/no inhibition and predicted the inhibition status of the test samples (studies 3 and 4) using the predict() function. The prediction area was visualized with a colored background on the sample plot, as described in [28]. Code and functions used for data analysis are described in [19, 28] and available at http://mixomics.org/ and https://gitlab.irstea.fr/olivier.chapleur/mint-bioindicators/.

## 3 Results

### 3.1 Integration of independent studies to identify microbial bioindicators

#### 3.1.1 Inhibition status classification of digesters according to methane production performance

A total of 81 samples were selected from studies 1 and 2. These samples were collected in 35 distinct digesters. Prior to identifying potential bioindicators characteristic of both type of inhibition, inhibition status of each digester (non-inhibited, inhibited by phenol or inhibited by ammonia) has been characterized. For this purpose, maximum CH_4_ production rate (mL CH_4_/day) was chosen as the most informative performance indicator. These values were calculated for each digester using Grofit package of R CRAN software (version 3.1.2) in both previous studies[20, 21]. To integrate both studies, we decided that samples were inhibited as soon as maximum CH_4_ production rate decreased by more than 20% compared to control and not inhibited if CH_4_ production rate decreased by less than 10%. Figure 1 presents boxplots describing the distribution of the relative decrease of maximum CH_4_ production rate of each digester according to their inhibition status.

**Figure 1:**
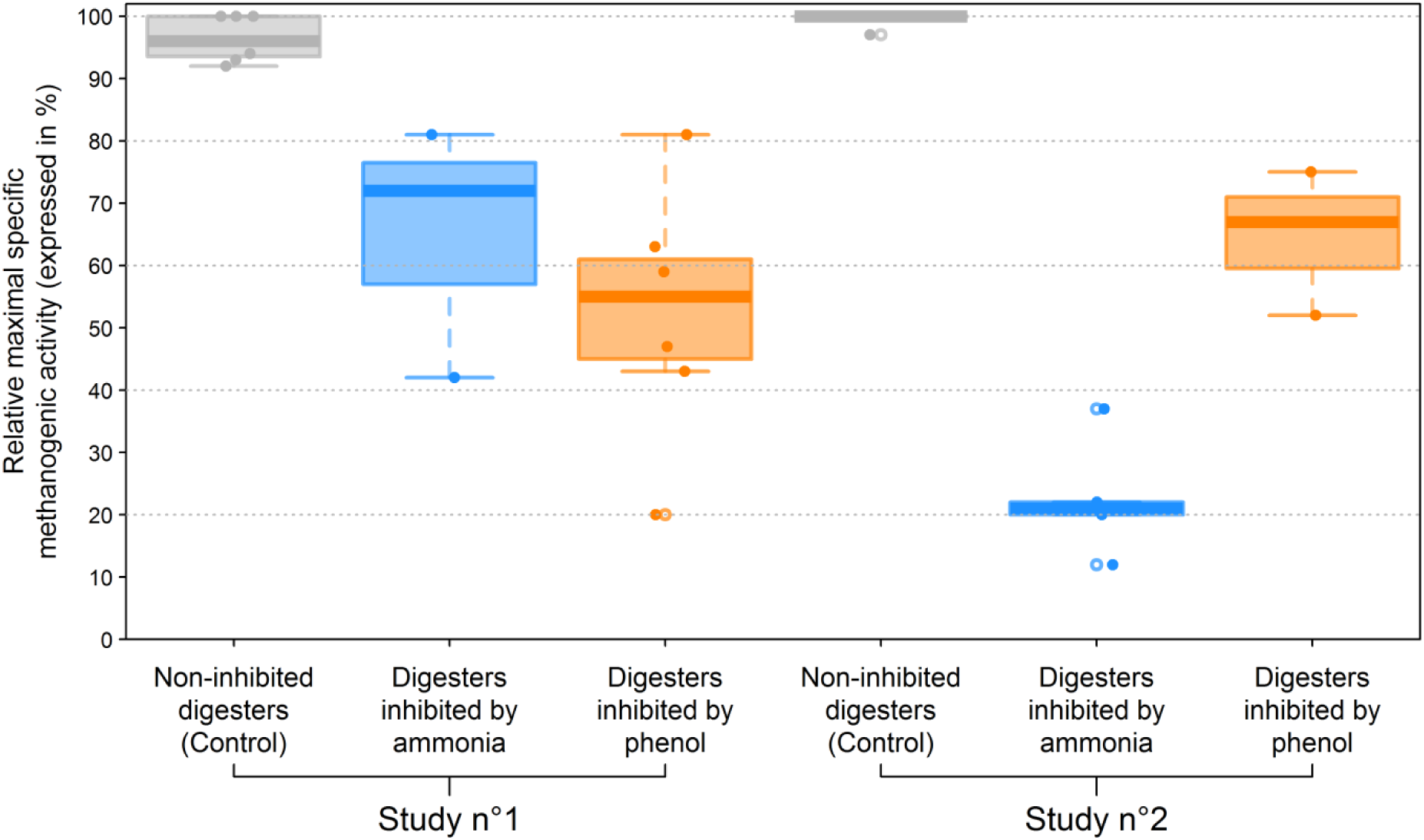
Relative maximal specific methanogenic activity in the digesters of studies 1 and 2. The different boxplots correspond to the different groups of digesters in each study. Relative maximal specific methanogenic activity was calculated as the ratio of maximal specific methanogenic activity in a digester divided by the maximal specific methanogenic activity in controls without inhibitor.

According to this threshold, we determined that 29 samples were non-inhibited whereas 24 samples were inhibited by phenol and 28 samples by ammonia. In study 1, samples were non-inhibited as soon as initial inhibitor concentration remained lower than 0.1 g/L of phenol or 2.5 g/L of TAN. In digesters inhibited by ammonia and phenol a decrease by respectively 20 to 60% and 20 to 80% of methanogenic activity was observed. In study 2, regardless support addition, all digesters facing 19g/L of TAN were considered as inhibited (decrease of methanogenic activity by 60 to 90%). In presence of 1.5g/L of phenol, only digesters supplemented with activated carbons were considered as non-inhibited.

#### 3.1.2 MINT modelling accounts for study effect

Considering, the inhibition status classification according to methane production performance, PCA was performed on the data (Fig. 2A), for a first exploration of the major sources of variation in the data. Sample distribution highlighted a strong study effect. Samples on the left part of the factorial plane were related to study 1 conducted with the inoculum A while samples collected during study 2 conducted with inoculum B were on the right side of the factorial plane.

**Figure 2:**
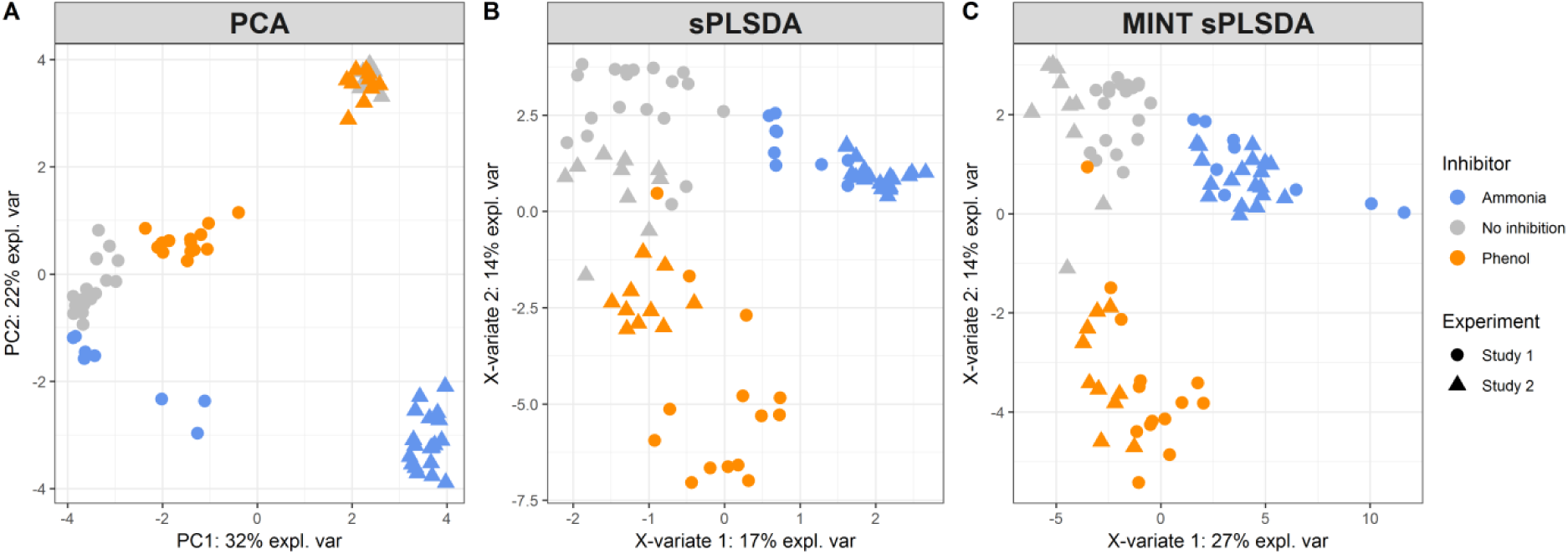
(A) Principal Component Analyses (PCA), (B) Sparse Partial Least Squares Discriminant Analysis, (C) Multivariate Integrative Sparse Partial Least Squares Discriminant Analysis of OTUs distribution in samples from studies 1 and 2, inhibited by phenol, ammonia or not inhibited. On the factorial maps each sample is represented with a coloured marker. The colour scale represents the type of inhibitor. The type of marker represents the study. OTU data was generated by 16S rRNA gene sequencing.

However, a clear influence of the type of inhibition on microbial community could still be observed. For both studies, ecosystems facing ammonia inhibition were strongly discriminated from samples that were non-inhibited or inhibited by phenol. Similarly, within the study 1 conducted without support media, samples collected from batch digester inhibited by phenol were separated from non-inhibited samples.

A supervised PLS-DA model was then fitted on the data. Sparse version of the method was applied to feature selection and identification of discriminative OTUs that best described the difference between groups of samples. In order to conduct sPLS-DA, parameters such as the number of components, and the number of OTUs to select must be specified. We chose these parameters based on the classification performance of sPLS-DA using cross-validation. Thirty-nine OTUs were thus selected by sPLS-DA and achieved a balanced error rate to 7.0% (2 components). Samples distribution based on the first two components is presented on Fig. 2B. sPLS-DA model enabled to mitigate the study effect compared to the unsupervised PCA. However, within each condition, the study effect was still present: each sample collected in Study 1 was clearly separated from the ones collected in Study 2.

In order to counteract this bias, we applied MINT that combines independent studies measured on the same OTU predictors and identifies reproducible bioindicator signatures across heterogeneous studies. As described above for sPLS-DA, we chose the optimal number of components and number of OTUs to select based on cross-validation, resulting in 45 OTUs and achieved a balanced error rate to 9.2% (2 components) (supplementary material table S2). Samples representation from MINT is presented in Fig. 2C. It evidenced that the study effect was accounted for, with the strongest separation observed according to inhibiting condition rather than studies. The classification error rate of the final MINT model was 9.2% confirming the good performance of MINT to classify our samples and identify a microbial signature.

#### 3.1.3 Analysis of microbial community

Microbial signatures identified with MINT were output in a clustered image maps (81 samples and 45 OTUs) in Fig. 3. This representation confirmed that, based on their microbial community composition, samples could be grouped by inhibition type (non-inhibited samples, samples inhibited by ammonia and samples inhibited by phenol). Moreover, the 45 OTUs selected by MINT were clustered into five different groups. The first group (Group A) was composed of 9 OTUs which were specifically correlated to digesters inhibited by phenol. Similarly, a second group of 17 OTUs (Group E) was associated to samples inhibited by ammonia. In group D, 6 OTUs were characteristic of both inhibitory conditions (phenol and ammonia). Group C included of 6 OTUs characterizing non-inhibited ecosystems while Group B was composed of 7 OTUs not recovered under ammonia inhibition. Interestingly, 6 of the 7 OTUs recovered in Group B were found in samples where phenol degradation was advanced. Consequently, the presence of these OTUs in this group may be explained by the variability of the inhibitory pressure throughout the incubation because of phenol degradation, and thus to their resilience capacity after phenol inhibition. Our following results are reported at the genus level, which was the most precise taxonomic level we could obtain with 16S rRNA sequencing.

**Figure 3:**
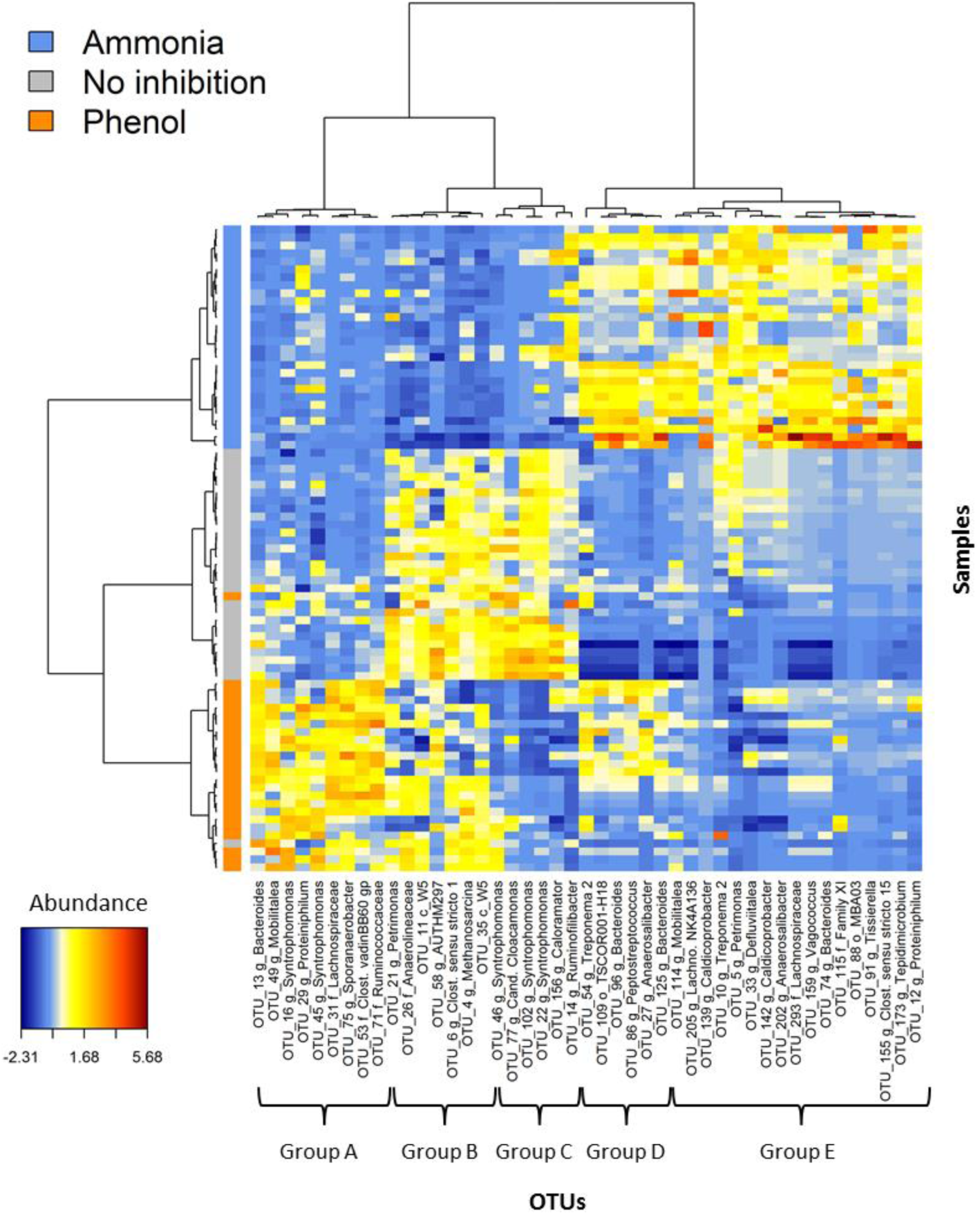
Heatmap of the most discriminant OTUs. Heatmap was built after selection of the most discriminant OTUs with Multivariate Integrative Sparse Partial Least Squares Discriminant Analysis of all OTUs generated by 16S rRNA gene sequencing for the different samples of studies 1 and 2, inhibited by phenol, ammonia or not inhibited. Name of the OTUs is indicated at the bottom. The color scale on the left represents the type of inhibitor. The color key of the heatmap shows the abundance of the OTUs after CLR transformation (from blue = low abundance to brown red = high abundance).

##### Genera correlated to non-inhibitory conditions

The only discriminative archaeal OTU evidenced by the model, belonging to *Methanosarcina* genus, was negatively correlated to ammonia inhibition. Its belonging to Group B indicated that this genus was also inhibited by phenol but could still grow once phenol was degraded, confirming that this genus is characteristic of non-inhibitory conditions. *Methanosarcina* genus is known to play a key role in anaerobic digestion of many feedstocks [30, 31]. However, although its robustness has been evidenced during different disruption situations [32, 33], it appeared that a drop of its relative abundance can reveal the presence of inhibitors such as ammonia or phenol in digesters. OTU_156 and OTU_6 assigned to *Clostridiaceae* 1 family and more precisely to *Caloramator* and *Clostridium butyricum*, strictly anaerobic acetogenic species described as butyrate producer [34] were also correlated to non-inhibitory conditions. This result confirmed the observation that was made for studies 1 and 2 during which butyrate production was inhibited by phenol and ammonia [12, 15]. While butyrate, acetate and propionate are regularly mentioned as indicators of process imbalance [35], our study tends to indicate that early variations in butyrate producers abundances could also be used as a bioindicator of inhibition. Interestingly, a single member of *Chloroflexi* phylum (OTU_26) kept for our model was also greatly associated to digesters for which phenol degradation had occurred. It was only assigned to the family *Anaerolinaceae*, which is widespread in full-scale digesters [36]. Its filamentous form was suggested to favor synergistic relationship with archaeal populations such as *Methanosaeta* also known to be inhibited by phenol and ammonia [37]. *Cloacimonetes* phylu*m* (formerly known as WWE1) was also specifically and strongly correlated to digesters where phenol degradation was advanced. It was represented by three distinct OTUs (OTU_11, OTU_35 and OTU_77). Only OTU_77 could be assigned at the genus level as *Cloacomonas*. As *Anaerolinaceae*, this genus was recovered in various studies analyzing methanogenic sludge which suggested it was a syntrophic bacterium capable of degrading propionate, amino acids and cellulose [38, 39]. Another study indicated that this genus was sensitive to different inhibitions and notably to the antibiotic monensin which is released in cow manure and recovered in anaerobic digester [40]. The last phylum that was specifically correlated to digesters where phenol was partly degraded was *Thermotogaceae* with a single OTU (OTU_58) assigned to *Petrotogaceae*, which is known to be involved in phenol degradation [41].

##### Genera correlated to a single type of inhibition

The model permitted the identification of genera specifically related to each type of inhibition. Interestingly, among the 17 OTUs that were correlated with ammonia inhibition, 9 of them belonged to 8 genera exclusively recovered in digesters inhibited by ammonia. They were assigned *Caldicoprobacter, Clostridium sensu stricto 15, Defluviitalea, Tepidimicrobium, Vagococcus, Tissierella, Lachnospiraceae NK4136* group and to an unknown genus of the *Family XI.* Among them, recent studies found that *Caldicoprobacter* populations were dominant in reactors exposed to high levels of ammonia [15, 42]. Similarly, Rui *et al*., evidenced that *Tissierella* genus was positively correlated to high concentration of TAN [43] while *Defluviitalea* were also dominant in digesters treating animal manure [44].

For the phenol, three OTUs belonging to *Clostridiales* order and assigned to genus *Sporoanaerobacter* (*Family XI*), to an unknown genus of the *vadinBB60* family and to an unassigned genus of the *Ruminococcaceae* family were evidenced. However, these cosmopolitan families are widely represented in anaerobic digesters.

On the other hand, we observed that some genera were similarly associated to digesters inhibited by phenol and to non-inhibited digesters suggesting that they were specifically sensitive to ammonia inhibition but not to phenol inhibition. It was notably the case for OTUs belonging to *Syntrophomonadaceae*, one of the major families of AD ecosystems. It consisted of five OTUs all assigned to *Syntrophomonas* genus but to unknown species. According to the model, three of them were related to non-inhibited digesters while two others were linked to phenol inhibition. *Syntrophomonas* are obligate anaerobic and syntrophic bacteria, which have the ability to oxidize saturated fatty acids, which is expected to enhance VFAs consumption. These functions are crucial in AD process. Thus, this result tended to confirm that under inhibitory conditions, reorganizations within the *Syntrophomonas* populations which carry these functions occur, thus confirming that the plasticity of the ecosystem is directly responsible for its resistance and resilience capacities [45]. Notably, the high functional redundancy among *Syntrophomonas* species seemed to allow the preservation of global metabolic chain and methane production.

##### Genera correlated to both inhibitions

A few bioindicators were also associated to both types of inhibition. It was notably the case for the four OTUs belonging to genus *Bacteroidetes*, as well as for both couples of OTUs belonging to family *Porphyromonadaceae* and assigned to *Petrimonas* and *Proteiniphilum*. These three genera have been acknowledged to play an important role in degrading complex carbohydrates and proteins and catalyzing the production of VFAs and CO_2_. Furthermore, the maintenance of important percentages of *Bacteroidetes* within a digester has already been suggested to be responsible for the ability of the anaerobic microbiota to counteract disturbances such as shock loadings [38]. Similarly, within *Clostridiales* order, genera *Anaerosalibacter, Mobilitalea, Peptostreptococcus* and two unknown genera of *Lachnospiraceae* family were correlated to both inhibitions. *Lachnospiraceae* and particularly to genus *Mobilitalea* as well as *Peptostreptococcus* and *Anaerosalibacter* were described as resistant to phenol and ammonia inhibition and have been suggested to play important roles in protein hydrolysis (Biddle et al.,2013). They were also reported to hydrolyze a variety of polysaccharides by different mechanisms.

##### *Clostridiales* are key bioindicators of inhibition

Interestingly, 25 out of the 45 discriminant OTUs selected in our model belonged to *Clostridiales* order. Among these 25 OTUs, 20 were related to inhibited ecosystems, which tended to indicate that inhibitory pressure by phenol or by ammonia would preferentially select more resistant bacteria affiliated to this order. A dominance of the *Clostridiales* order has been reported by many studies at suboptimal conditions for methanogenesis (increased ammonia or salt concentrations) [46, 47]. Moreover, the importance of this class is regularly mentioned as crucial in AD process [48]. However, the majority of the OTUs belonging to this class could not be specifically correlated with only one type of inhibition. It could be hypothesized that it may be due to the high functional redundancy within these genera and diverse populations as mentioned previously. This limitation was also observed for other phyla and notably for *Bacteroidetes* and *Spirochaetae*. Furthermore, the lack of sequencing precision associated to the short length of 16S regions amplified prevented the affiliation at taxonomic rank such as species or subspecies. Despite the fact that we could not conclude about the correlation between these OTUs and the type of inhibition, the model still highlighted that the emergence of these genera were associated to a selection pressure caused by phenol or ammonia. Notably, it confirmed the negative correlation observed by Heyer *et al*., between *Methanosarcinales*, which is a marker of steady state and bacterial orders such as *Clostridiales* and *Spirochaetales* [49]. It also reinforced the link found by Lee *et al*. between *Spirochaetales* and inhibited ecosystems [50].

### 3.2 A predictive model for ammonia inhibition validated in two external studies

Samples from external studies 3 and 4 were analyzed with distinct sequencing techniques. Thus, different bioinformatic treatments were applied to raw sequences of each dataset due to differences in sequencing primers and targeted regions. OTUs were subsequently aggregated at the genus level (called ‘clusters’) in order to merge the three datasets. Since both external studied were focused on inhibition by ammonia only, we trained a new MINT model from our in-house samples where we removed the phenol condition. A total of 55 samples collected during studies 1 and 2 were retained, including 29 samples considered as non-inhibited and 26 as inhibited by ammonia.

#### 3.2.1 Inhibition status prediction of external samples

In order to be consistent with the experimental strategy of the authors, samples from Lü et al., study were categorized into four groups depending on the sampling time and on the inhibitory pressure. Samples collected from digesters non-inhibited with ammonia were gathered in the “No inhibition” group. Samples inhibited with 3g/L of TAN were clustered in the “Ammonia moderate concentration” group. Samples inhibited with 7g/L of TAN were divided into two groups: “Ammonia inhibition, early days” for samples collected at the end of the lag phase and “Ammonia inhibition, final days” for samples collected near the end of the methane-production phase. Similarly, samples from Peng et al., study were categorized into four groups depending on the operational time when they were collected. Samples from day 0 to day 127, where methane yield remained stable circa 0.5 mL CH_4_/g VS, were clustered in the “No inhibition” group. Samples from day 139 to day 152 were clustered into the “Ammonia inhibition start” group. During this phase, methane yield began to strongly fluctuate and slightly decrease down to 0.4 mL CH_4_/g VS. Samples from day 172 to day 212 during which methane yield dropped down to 0.25 mL CH_4_/g VS, were clustered in the “Ammonia inhibition” group. Both samples from day 223 to day 232 were clustered in the “Ammonia inhibition decrease” group where biogas production restarted while methane yield remained below 0.25 mL CH_4_/g VS.

#### 3.2.2 Ammonia inhibition model

As previously, PCA and sPLSDA were performed on the data (Fig. 4A and Fig. 4B). An optimal number of 17 clusters were selected by the model, leading to a balanced error rate of 1.7% for sPLD-DA Sample distribution before applying MINT confirmed the strong study effect. The efficiency of MINT to discriminate the in-house samples into inhibition / no-inhibition conditions is depicted in Fig. 4C for the first two MINT components, with the first (horizontal) component highly discriminative of the inhibitory status of the digester. Yet, three of the inhibited samples were located on the left hand side of the plot. They corresponded to the most inhibited samples of study 1 with the highest concentration of TAN (25 g/L).

**Figure 4:**
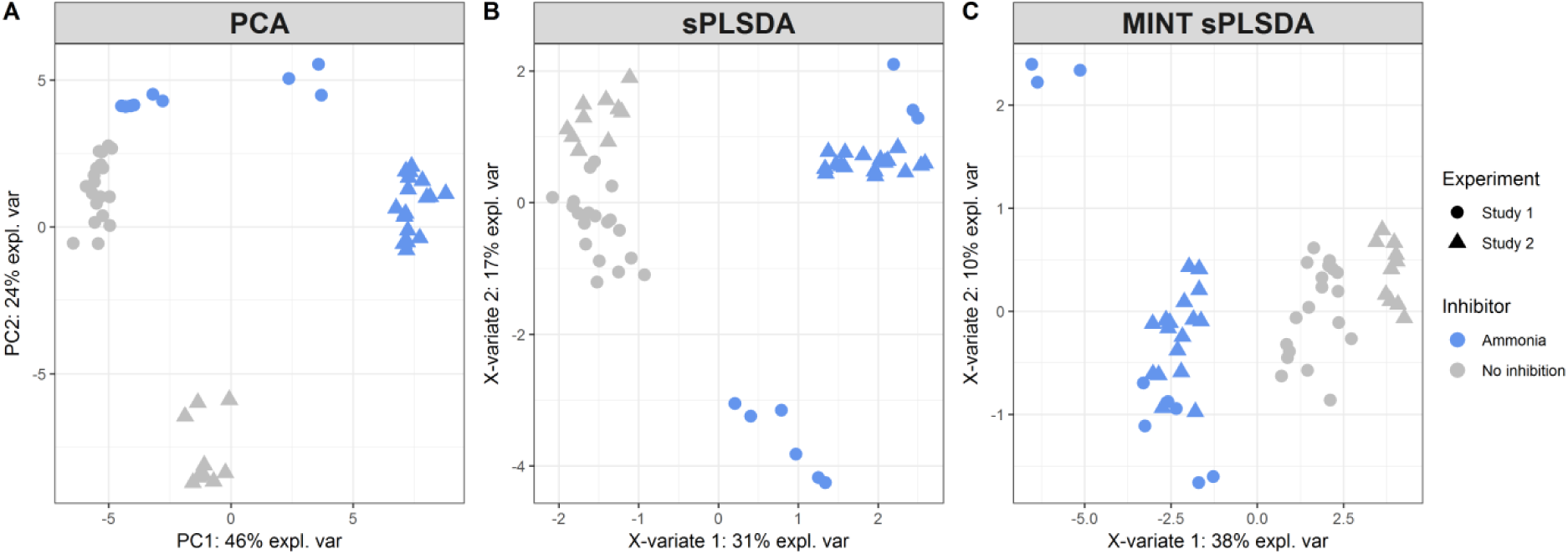
(A) Principal Component Analyses (PCA), (B) Sparse Partial Least Squares Discriminant Analysis, (C) Multivariate Integrative Sparse Partial Least Squares Discriminant Analysis of taxonomic distribution (genus level) in samples from studies 1 and 2, inhibited by ammonia or not inhibited. On the factorial maps each sample is represented with a coloured marker. The colour scale represents the type of inhibitors. The type of marker represents the study. Taxonomic data was generated by 16S rRNA gene sequencing and aggregation of the data at the genus level.

The OTUs selected in this second MINT model (supplementary material table S3) were consistent with the observation made with the first model in our previous section (Figure 5). Among them, 10 clusters were positively correlated to non-inhibited digesters. Notably, it confirmed that *Clostridium butyricum* (Cluster_6) could be considered as a robust bioindicator of non-inhibitory conditions. Furthermore, clusters assigned to *Cloacimonetes* phylum (Cluster_11, Cluster_43 and Cluster_49) and particularly to genus *Cloacomonas* clearly related to non-inhibited digesters were re-evidenced. Nevertheless, due to the novelty of this phylum, missing reference genomes most probably hampered a deeper taxonomic classification of *Cloacimonetes* species. The ecological function of *Cloacimonetes* is thus not established, but this group is suggested to be only present in mesophilic conditions, involved in amino acid fermentation, syntrophic propionate oxidation and extracellular hydrolysis [51]. It was also evidenced that its abundance decreased with increasing ammonia levels [52]. We also confirmed that *Syntrophomonas* (Cluster_17) was more represented in absence of ammonia. Moreover, the second model also revealed three new genera considered as discriminant of non-inhibited digesters conditions. These genera were assigned to two families belonging to *Clostridiales* order: *Ruminococcaceae* (Cluster_34 and Cluster_69), and *Christensenellaceae* (Cluster_42). Nevertheless, genera belonging to *Clostridiales* order are known to be involved in various metabolic activities which prevent us from elucidating their specific role under non-inhibiting conditions.

**Figure 5:**
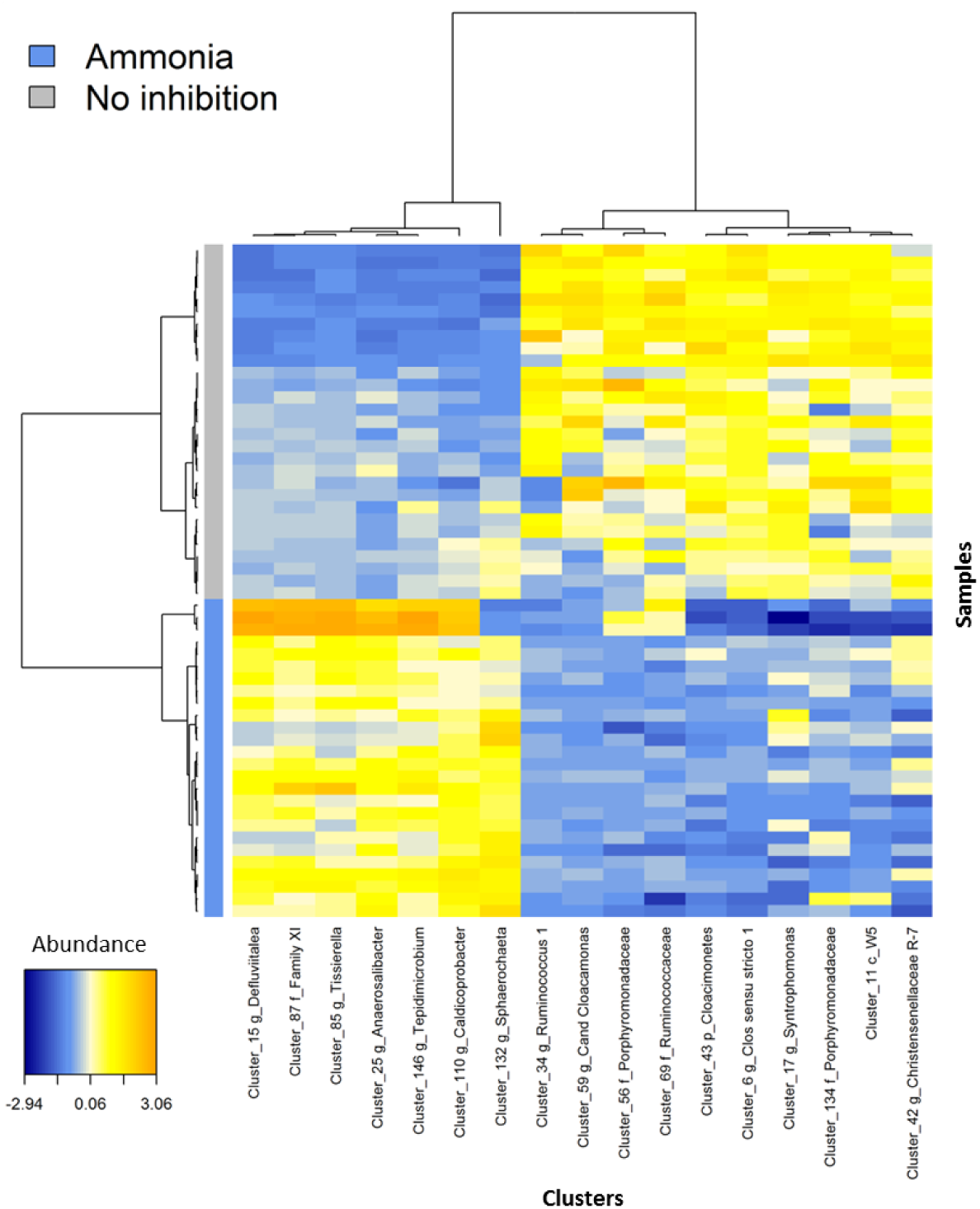
Heatmap of the most discriminant clusters of OTUs. Heatmap was built after selection of the most discriminant clusters with Multivariate Integrative Sparse Partial Least Squares Discriminant Analysis of all clusters generated after OTUs aggregation at the genus level for the different samples of studies 1 and 2, inhibited ammonia or not inhibsited. Name of the clusters is indicated at the bottom. The color scale on the left represents the type of inhibitor. The color key of the heatmap shows the abundance of the clusters after CLR transformation (from blue = low abundance to orange = high abundance).

On the other hand, seven clusters were strongly linked to ammonia inhibition. Six of them belonged to *Clostridiales* order and notably to genera that were previously identified in the first model as specific markers of this type of inhibition (*Caldicoprobacter, Defluviitalea, Anaerosalibacter, Tepidimicrobium* and *Tissierella*). Another unknown genus belonging to *Family XI* was also discriminant confirming the great resistance of this family to ammonia inhibitory pressure. The last OTU was assigned to *Sphaerochaeta*, which belongs to *Spirochaetales* order. This order was associated to both types of inhibitions in the previous model. Interestingly, the heatmap presented in Fig. 5 emphasized that six of these seven clusters were significantly more abundant in samples collected in digesters inhibited with the highest ammonia concentration, thus reinforcing the robustness of these bioindicators.

The bioindicators highlighted by the second model were consistent with those evidenced by the model integrating phenol inhibition, thus confirming the robustness of this statistical analysis.

#### 3.2.3 Prediction of the inhibitory status of samples analyzed by external studies

This second model was built to predict the inhibitory status of samples collected during two external studies (studies 3 and 4). The prediction results are indicated in Table 2. In order to visualize external samples distribution in the model, Fig. 6 presents the test samples from studies 3 and 4 projected on the two first components of the trained model, as well as prediction areas [28].

**Table 2:** Samples of studies 3 and 4, observed inhibition status based on the original paper and predictions of the model

| Study | Sample accession number | Sample number | Inhibition status (based on the paper) | Prediction |
| --- | --- | --- | --- | --- |
| **Study 3** | SRR1507064_1 | 1 | No inhibition | No inhibition |
|  | SRR1507067_1 | 2 | No inhibition | No inhibition |
|  | SRR1507069_1 | 3 | No inhibition | No inhibition |
|  | SRR1507183_1 | 4 | No inhibition | No inhibition |
|  | SRR1507184_1 | 5 | No inhibition | No inhibition |
|  | SRR1507185_1 | 6 | No inhibition | No inhibition |
|  | SRR1507186_1 | 7 | No inhibition | No inhibition |
|  | SRR1507187_1 | 8 | No inhibition | No inhibition |
|  | SRR1507188_1 | 9 | No inhibition | No inhibition |
|  | SRR1507189_1 | 10 | No inhibition | No inhibition |
|  | **SRR1507190_1** | **11** | **No inhibition** | **Ammonia** |
|  | SRR1507191_1 | 12 | No inhibition | No inhibition |
|  | SRR2296764_1 | 13 | Ammonia moderate concentration | No inhibition |
|  | SRR2296765_1 | 14 | Ammonia moderate concentration | Ammonia |
|  | SRR2296766_1 | 15 | Ammonia moderate concentration | No inhibition |
|  | SRR2296767_1 | 16 | Ammonia moderate concentration | No inhibition |
|  | SRR2296768_1 | 17 | Ammonia inhibition, final days | Ammonia |
|  | SRR2296769_1 | 18 | Ammonia inhibition, early days | Ammonia |
|  | SRR2296770_1 | 19 | Ammonia inhibition, early days | Ammonia |
|  | SRR2296771_1 | 20 | Ammonia inhibition, early days | Ammonia |
|  | SRR2296772_1 | 21 | Ammonia inhibition, early days | Ammonia |
|  | SRR2296773_1 | 22 | Ammonia inhibition, final days | Ammonia |
|  | SRR2296774_1 | 23 | Ammonia inhibition, final days | Ammonia |
|  | SRR2296775_1 | 24 | Ammonia inhibition, final days | Ammonia |
|  | SRR2296776_1 | 25 | Ammonia inhibition, early days | Ammonia |
|  | SRR2296777_1 | 26 | Ammonia inhibition, early days | No inhibition |
|  | SRR2296778_1 | 27 | Ammonia inhibition, early days | No inhibition |
|  | SRR2296779_1 | 28 | Ammonia inhibition, final days | Ammonia |
|  | SRR2296780_1 | 29 | Ammonia inhibition, final days | Ammonia |
|  | SRR2296781_1 | 30 | Ammonia inhibition, final days | Ammonia |
|  | SRR2296782_1 | 31 | Ammonia inhibition, early days | No inhibition |
|  | SRR2296783_1 | 32 | Ammonia inhibition, early days | No inhibition |
|  | SRR2296784_1 | 33 | Ammonia inhibition, early days | Ammonia |
|  | SRR2296785_1 | 34 | Ammonia inhibition, final days | Ammonia |
|  | SRR2296786_1 | 35 | Ammonia inhibition, final days | Ammonia |
|  | SRR2296787_1 | 36 | Ammonia inhibition, final days | Ammonia |
| **Study 4** | SRR3629052 | 37 | No inhibition | No inhibition |
|  | SRR3629054 | 38 | No inhibition | No inhibition |
|  | **SRR3629058** | **39** | **No inhibition** | **Ammonia** |
|  | SRR3629059 | 40 | Ammonia inhibition start | No inhibition |
|  | SRR3629149 | 41 | Ammonia inhibition start | No inhibition |
|  | SRR3629150 | 42 | Ammonia inhibition | Ammonia |
|  | SRR3629151 | 43 | Ammonia inhibition | Ammonia |
|  | SRR3629152 | 44 | Ammonia inhibition | Ammonia |
|  | SRR3629153 | 45 | Ammonia inhibition decrease | Ammonia |
|  | SRR3629154 | 46 | Ammonia inhibition decrease | No inhibition |

**Figure 6:**
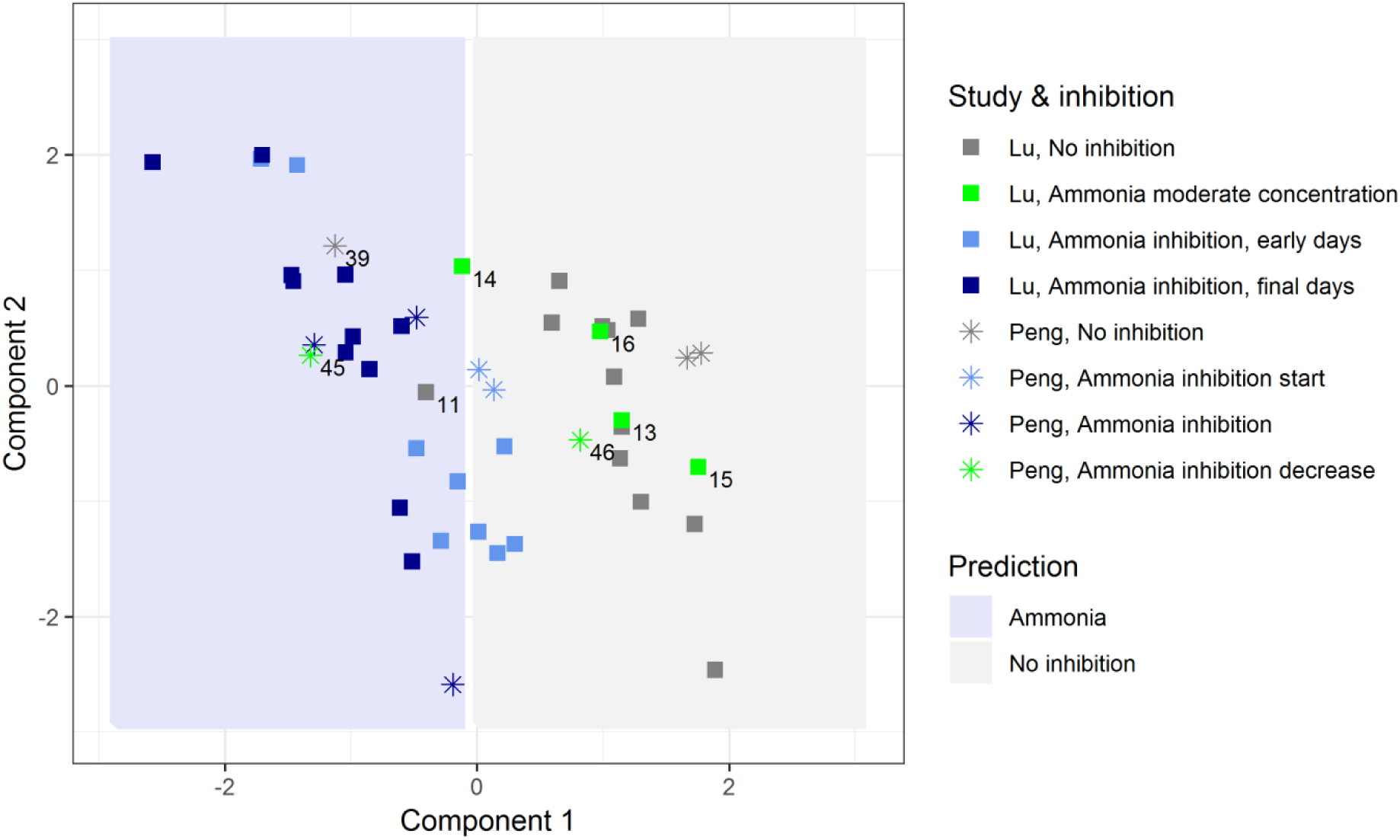
Projection of samples from studies 3 and 4 in the factorial plan determined after Multivariate Integrative Sparse Partial Least Squares Discriminant Analysis of samples from studies 1 and 2. Each sample from studies 3 and 4 is represented by a marker. Type of marker indicates the study. Color of the marker indicates the inhibition status in the reactor where the sample was taken. A prediction area, based on studies 1 and 2 was calculated and is plotted on the graph. The different figures indicate remarkable samples.

As expected, inhibited samples were separated against the non-inhibited samples on the first component. Two samples (11 and 39) were mispredicted as ‘inhibited’. Samples of reactors that just started inhibition (“early days” in Lü et al. and ‘start of the inhibition’ in Peng et al.) were mostly predicted at an intermediary position and predicted as either inhibited or non-inhibited. We hypothesized that microbial community had started to change but was not yet totally characteristic of inhibited reactors. Interestingly, sample 45 (Peng et al., day 223, just after inhibitory pressure was lowered) was predicted as inhibited while sample 46 (Peng et al. day 232, several days after inhibitory pressure was lowered) was predicted as non-inhibited. This result may illustrate the progressive resilience of the microbial community after the inhibition. Sample 14 (moderate ammonia, final days, no addition of activated charcoal) was predicted as inhibited while samples 13, 15, 16 (moderate ammonia, but early days or addition of activated charcoal) were predicted as non-inhibited, in agreement with the conclusions of the authors. Finally, taking into account all the samples from digesters clearly non-inhibited (15) or clearly inhibited by ammonia (13) we estimated that the model predicted the inhibitory status of external samples with an accuracy of 93% as only two samples were incorrectly predicted.

## 4 Discussion

16S rRNA gene sequencing have revolutionized environmental biotechnology research and helped to progressively unravel the complexity of AD inhibition, but we still need to improve the resilience of AD systems and promote its implementation at a larger scale to avoid system failure. Our study focused on identifying robust microbial indicators in two in-house studies that, in combination in our multivariate model, were highly predictive of inhibition status in two external studies.

Despite the complexity and the functional redundancy of the microbial community within digesters, our model revealed the feasibility of detecting key indicators evidencing the state of the AD process whilst addressing the challenge of study-specific effects. The microbial indicators we identified were separate from cosmopolitan OTUs that tended to co-occur in all conditions. Thus, these indicators can be considered as bioindicators to announce early signs of process dysfunction in anaerobic digesters. Our study emphasized on the benefit of using multivariate models to predict the inhibitory status of external studies based on a microbial signature identified in a training set (here our two-in-house studies), despite differences in sequencing primers and targeted regions.

When successful, integrative multi-studies analysis not only allows to increase in sample size and statistical power, but also to share data across research communities and re- use existing data deposited in public databases with the common goal of identifying reproducible microbial signatures. Our results are encouraging as they suggest potential applications for a wide diversity of AD reactors and biotechnologies, and thus pave the way for digester management given the uncertainty related to inhibition thresholds in individual reactors. As an increasing number of 16S sequences databases are available to build models predicting different types of digester inhibitions for different kind of digesters operating in various conditions, such results will allow to optimize the prediction quality by recursively updating monitoring model. We anticipate that a single metagenetic sequencing will reduce the number and the complexity of analyses targeting different inhibitors, as bioindicators identified could be correlated to different types of inhibitions. Moreover, miniaturization and portability of sequencers will soon allow high frequency on-line measurements of microbial dynamics, involving low workload, and limited sampling issues [53]

The next step will be further our investigations to define the role of these microorganisms in AD process. As 16S reference databases are currently still incomplete, taxonomic assignment at the genus level results in a substantial lack of data interpretation. The use of shotgun metagenomic and metatranscriptomic tools should also be useful to get different type of information such as functional bioindicators.

## 5 Conclusions

- A robust microbial signature of AD inhibition by ammonia and phenol was identified by integrating independent16S metabarcoding studies with statistical approaches.
- These biomarkers were successfully used to predict the inhibition in independent digesters with a multivariate model.
- Our approach can be generalized to other inhibitors and other studies to build robust models of AD inhibition useful for an improved management of anaerobic digesters.

## References

1. Scarlat N., Dallemand J.-F., and Fahl F. Biogas: Developments and perspectives in Europe. Renewable Energy. 2018; 129: 457–472.

2. Rajagopal R., Masse D.I., and Singh G. A critical review on inhibition of anaerobic digestion process by excess ammonia. Bioresour Technol. 2013; 143: 632–41.

3. Gonzalez-Gil L., Mauricio-Iglesias M., Serrano D., Lema J.M., and Carballa M. Role of methanogenesis on the biotransformation of organic micropollutants during anaerobic digestion. Sci Total Environ. 2018; 622-623: 459–466.

4. Yenigün O. and Demirel B. Ammonia inhibition in anaerobic digestion: A review. Process Biochemistry. 2013; 48: 901–911.

5. Rosenkranz F., Cabrol L., Carballa M., Donoso-Bravo A., Cruz L., Ruiz-Filippi G., et al. Relationship between phenol degradation efficiency and microbial community structure in an anaerobic SBR. Water Res. 2013; 47: 6739–49.

6. Veeresh G.S., Kumar P., and Mehrotra I. Treatment of phenol and cresols in upflow anaerobic sludge blanket (UASB) process: a review. Water Res. 2005; 39: 154–70.

7. Appels L., Lauwers J., Degrève J., Helsen L., Lievens B., Willems K., et al. Anaerobic digestion in global bio-energy production: Potential and research challenges. Renewable and Sustainable Energy Reviews. 2011; 15: 4295–4301.

8. Li Y., Chen Y., and Wu J. Enhancement of methane production in anaerobic digestion process: A review. Applied Energy. 2019; 240: 120–137.

9. Calusinska M., Goux X., Fossepre M., Muller E.E.L., Wilmes P., and Delfosse P. A year of monitoring 20 mesophilic full-scale bioreactors reveals the existence of stable but different core microbiomes in bio-waste and wastewater anaerobic digestion systems. Biotechnol Biofuels. 2018; 11: 196.

10. Li L., He Q., Ma Y., Wang X., and Peng X. Dynamics of microbial community in a mesophilic anaerobic digester treating food waste: Relationship between community structure and process stability. Bioresource Technology. 2015; 189: 113–120.

11. Wang P., Wang H., Qiu Y., Ren L., and Jiang B. Microbial characteristics in anaerobic digestion process of food waste for methane production–A review. Bioresource Technology. 2018; 248: 29–36.

12. Poirier S., Bize A., Bureau C., Bouchez T., and Chapleur O. Community shifts within anaerobic digestion microbiota facing phenol inhibition: Towards early warning microbial indicators? Water Res. 2016; 100: 296–305.

13. Chapleur O., Madigou C., Civade R., Rodolphe Y., Mazéas L., and Bouchez T. Increasing concentrations of phenol progressively affect anaerobic digestion of cellulose and associated microbial communities. Biodegradation. 2016; 27: 15–27.

14. Madigou C., Poirier S., Bureau C., and Chapleur O. Acclimation strategy to increase phenol tolerance of an anaerobic microbiota. Bioresour Technol. 2016; 216: 77–86.

15. Poirier S., Desmond-Le Quemener E., Madigou C., Bouchez T., and Chapleur O. Anaerobic digestion of biowaste under extreme ammonia concentration: Identification of key microbial phylotypes. Bioresour Technol. 2016; 207: 92–101.

16. Lu Y., Liaquat R., Astals S., Jensen P.D., Batstone D.J., and Tait S. Relationship between microbial community, operational factors and ammonia inhibition resilience in anaerobic digesters at low and moderate ammonia background concentrations. N Biotechnol. 2018; 44: 23–30.

17. Li J., Rui J., Yao M., Zhang S., Yan X., Wang Y., et al. Substrate Type and Free Ammonia Determine Bacterial Community Structure in Full-Scale Mesophilic Anaerobic Digesters Treating Cattle or Swine Manure. Frontiers in Microbiology. 2015; 6: 1337.

18. Werner J.J., Garcia M.L., Perkins S.D., Yarasheski K.E., Smith S.R., Muegge B.D., et al. Microbial Community Dynamics and Stability during an Ammonia-Induced Shift to Syntrophic Acetate Oxidation. Applied and Environmental Microbiology. 2014; 80: 3375–3383.

19. Rohart F., Eslami A., Matigian N., Bougeard S., and Lê Cao K.A. MINT: A multivariate integrative method to identify reproducible molecular signatures across independent experiments and platforms. BMC Bioinformatics. 2017; 18.

20. Poirier S. and Chapleur O. Influence of support media supplementation to reduce the inhibition of anaerobic digestion by phenol and ammonia: Effect on degradation performances and microbial dynamics. Data in Brief. 2018; 19: 1733–1754.

21. Poirier S. and Chapleur O. Inhibition of anaerobic digestion by phenol and ammonia: Effect on degradation performances and microbial dynamics. Data in Brief. 2018: 2235–2239.

22. Peng X., Zhang S., Li L., Zhao X., Ma Y., and Shi D. Long-term high-solids anaerobic digestion of food waste: Effects of ammonia on process performance and microbial community. Bioresour Technol. 2018; 262: 148–158.

23. Lü F., Luo C., Shao L., and He P. Biochar alleviates combined stress of ammonium and acids by firstly enriching Methanosaeta and then Methanosarcina. Water Research. 2016; 90: 34–43.

24. Zhang J., Kobert K., Flouri T., and Stamatakis A. PEAR: a fast and accurate Illumina Paired-End reAd mergeR. Bioinformatics. 2014; 30: 614–20.

25. Martin M. Cutadapt removes adapter sequences from high-throughput sequencing reads. 2011. 2011; 17: 3.

26. Escudié F., Auer L., Bernard M., Mariadassou M., Cauquil L., Vidal K., et al. FROGS: Find, Rapidly, OTUs with Galaxy Solution. Bioinformatics. 2018; 34: 1287–1294.

27. Camacho C., Coulouris G., Avagyan V., Ma N., Papadopoulos J., Bealer K., et al. BLAST+: architecture and applications. BMC Bioinformatics. 2009; 10: 421.

28. Rohart F., Gautier B., Singh A., and Lê Cao K.-A. mixOmics: An R package for ‘omics feature selection and multiple data integration. PLOS Computational Biology. 2017; 13: e1005752.

29. Lê Cao K.A., Costello M.E., Lakis V.A., Bartolo F., Chua X.Y., Brazeilles R., et al. MixMC: A multivariate statistical framework to gain insight into microbial communities. PLoS ONE. 2016; 11.

30. FitzGerald J.A., Allen E., Wall D.M., Jackson S.A., Murphy J.D., and Dobson A.D.W. Methanosarcina Play an Important Role in Anaerobic Co-Digestion of the Seaweed Ulva lactuca: Taxonomy and Predicted Metabolism of Functional Microbial Communities. PloS one. 2015; 10: e0142603–e0142603.

31. Vavilin V.A., Qu X., Mazeas L., Lemunier M., Duquennoi C., He P., et al. Methanosarcina as the dominant aceticlastic methanogens during mesophilic anaerobic digestion of putrescible waste. Antonie Van Leeuwenhoek. 2008; 94: 593–605.

32. De Vrieze J., Hennebel T., Boon N., and Verstraete W. Methanosarcina: the rediscovered methanogen for heavy duty biomethanation. Bioresour Technol. 2012; 112: 1–9.

33. Lins P., Reitschuler C., and Illmer P. Methanosarcina spp., the key to relieve the start-up of a thermophilic anaerobic digestion suffering from high acetic acid loads. Bioresour Technol. 2014; 152: 347–54.

34. Cassir N., Benamar S., and La Scola B. Clostridium butyricum: from beneficial to a new emerging pathogen. Clinical Microbiology and Infection. 2016; 22: 37–45.

35. Ahring B.K., Sandberg M., and Angelidaki I. Volatile fatty acids as indicators of process imbalance in anaerobic digestors. Applied Microbiology and Biotechnology. 1995; 43: 559–565.

36. Kirkegaard R.H., McIlroy S.J., Kristensen J.M., Nierychlo M., Karst S.M., Dueholm M.S., et al. Identifying the abundant and active microorganisms common to full scale anaerobic digesters. bioRxiv. 2017.

37. McIlroy S.J., Kirkegaard R.H., Dueholm M.S., Fernando E., Karst S.M., Albertsen M., et al. Culture-Independent Analyses Reveal Novel Anaerolineaceae as Abundant Primary Fermenters in Anaerobic Digesters Treating Waste Activated Sludge. Frontiers in microbiology. 2017; 8: 1134–1134.

38. Regueiro L., Spirito C.M., Usack J.G., Hospodsky D., Werner J.J., and Angenent L.T. Comparing the inhibitory thresholds of dairy manure co-digesters after prolonged acclimation periods: Part 2--correlations between microbiomes and environment. Water Res. 2015; 87: 458–66.

39. Limam R.D., Chouari R., Mazéas L., Wu T.D., Li T., Grossin-Debattista J., et al. Members of the uncultured bacterial candidate division WWE1 are implicated in anaerobic digestion of cellulose. MicrobiologyOpen. 2014; 3: 157–167.

40. Spirito C.M., Daly S.E., Werner J.J., and Angenent L.T. Redundancy in Anaerobic Digestion Microbiomes during Disturbances by the Antibiotic Monensin. Applied and Environmental Microbiology. 2018; 84.

41. Na J.-G., Lee M.-K., Yun Y.-M., Moon C., Kim M.-S., and Kim D.-H. Microbial community analysis of anaerobic granules in phenol-degrading UASB by next generation sequencing. Biochemical Engineering Journal. 2016; 112: 241–248.

42. Lv Z., Leite A.F., Harms H., Glaser K., Liebetrau J., Kleinsteuber S., et al. Microbial community shifts in biogas reactors upon complete or partial ammonia inhibition. Appl Microbiol Biotechnol. 2018.

43. Rui J., Li J., Zhang S., Yan X., Wang Y., and Li X. The core populations and co-occurrence patterns of prokaryotic communities in household biogas digesters. Biotechnology for biofuels. 2015; 8: 158–158.

44. Ma S., Huang Y., Wang C., Fan H., Dai L., Zhou Z., et al. Defluviitalea raffinosedens sp. nov., a thermophilic, anaerobic, saccharolytic bacterium isolated from an anaerobic batch digester treating animal manure and rice straw. Int J Syst Evol Microbiol. 2017; 67: 1607–1612.

45. Shade A., Peter H., Allison S.D., Baho D.L., Berga M., Burgmann H., et al. Fundamentals of microbial community resistance and resilience. Front Microbiol. 2012; 3: 417.

46. Alsouleman K., Linke B., Klang J., Klocke M., Krakat N., and Theuerl S. Reorganisation of a mesophilic biogas microbiome as response to a stepwise increase of ammonium nitrogen induced by poultry manure supply. Bioresour Technol. 2016; 208: 200–204.

47. De Vrieze J., Saunders A.M., He Y., Fang J., Nielsen P.H., Verstraete W., et al. Ammonia and temperature determine potential clustering in the anaerobic digestion microbiome. Water Research. 2015; 75: 312–323.

48. Joyce A., Ijaz U.Z., Nzeteu C., Vaughan A., Shirran S.L., Botting C.H., et al. Linking Microbial Community Structure and Function During the Acidified Anaerobic Digestion of Grass. Frontiers in Microbiology. 2018; 9.

49. Heyer R., Benndorf D., Kohrs F., De Vrieze J., Boon N., Hoffmann M., et al. Proteotyping of biogas plant microbiomes separates biogas plants according to process temperature and reactor type. Biotechnology for biofuels. 2016; 9: 155–155.

50. Lee S.-H., Park J.-H., Kang H.-J., Lee Y.H., Lee T.J., and Park H.-D. Distribution and abundance of Spirochaetes in full-scale anaerobic digesters. Bioresource Technology. 2013; 145: 25–32.

51. Muller B., Sun L., Westerholm M., and Schnurer A. Bacterial community composition and fhs profiles of low- and high-ammonia biogas digesters reveal novel syntrophic acetate-oxidising bacteria. Biotechnol Biofuels. 2016; 9: 48.

52. Westerholm M., Isaksson S., Karlsson Lindsjö O., and Schnürer A. Microbial community adaptability to altered temperature conditions determines the potential for process optimisation in biogas production. Applied Energy. 2018; 226: 838–848.

53. Shaffer L. Inner Workings: Portable DNA sequencer helps farmers stymie devastating viruses. Proceedings of the National Academy of Sciences. 2019; 116: 3351.

