## Supplemental Table S1, table S2, table S3 for "Integrating independent microbial studies to build predictive models of anaerobic digestion inhibition by ammonia and phenol"

**Table S1: list of samples included in the analysis**

| Study number | Sample name | Inhibitory status | Number of days at sampling |
|---|---|---|---|
| Study 1 | DNA.0P2T4 | No inhibition | 29 |
| | DNA.0P2T6 | No inhibition | 57 |
| | DNA.10P2T4 | No inhibition | 29 |
| | DNA.10P2T6 | No inhibition | 57 |
| | DNA.75P2T4 | Phenol inhibition | 29 |
| | DNA.75P2T6 | Phenol inhibition | 57 |
| | DNA.100P2T4 | Phenol inhibition | 29 |
| | DNA.100P2T6 | Phenol inhibition | 57 |
| | DNA.125P1T4 | Phenol inhibition | 29 |
| | DNA.125P1T6 | Phenol inhibition | 57 |
| | DNA.125P2T4 | Phenol inhibition | 29 |
| | DNA.125P2T6 | Phenol inhibition | 57 |
| | DNA.125P3T4 | Phenol inhibition | 29 |
| | DNA.125P3T6 | Phenol inhibition | 57 |
| | DNA.150P2T4 | Phenol inhibition | 29 |
| | DNA.150P2T6 | Phenol inhibition | 57 |
| | DNA.200P2T4 | Phenol inhibition | 29 |
| | DNA.200P2T6 | Phenol inhibition | 57 |
| | DNA.0N2T4 | No inhibition | 29 |
| | DNA.0N2T5 | No inhibition | 42 |
| | DNA.0N2T6 | No inhibition | 57 |
| | DNA.5N2T4 | No inhibition | 29 |
| | DNA.5N2T5 | No inhibition | 42 |
| | DNA.5N2T6 | No inhibition | 57 |
| | DNA.10N2T4 | No inhibition | 29 |
| | DNA.10N2T5 | No inhibition | 42 |
| | DNA.10N2T6 | No inhibition | 57 |
| | DNA.15N2T4 | No inhibition | 29 |
| | DNA.15N2T5 | No inhibition | 42 |
| | DNA.15N2T6 | No inhibition | 57 |
| | DNA.25N2T4 | No inhibition | 29 |
| | DNA.25N2T5 | No inhibition | 42 |
| | DNA.25N2T6 | No inhibition | 57 |
| | DNA.75N2T4 | Ammonia inhibition | 29 |
| | DNA.75N2T5 | Ammonia inhibition | 42 |
| | DNA.75N2T6 | Ammonia inhibition | 57 |
| | DNA.100N2T4 | Ammonia inhibition | 29 |
| | DNA.100N2T5 | Ammonia inhibition | 42 |
| | DNA.100N2T6 | Ammonia inhibition | 57 |
| | DNA.250N2T4 | Ammonia inhibition | 29 |
| | DNA.250N2T5 | Ammonia inhibition | 42 |
| | DNA.250N2T6 | Ammonia inhibition | 57 |

| | | | |
|---|---|---|---|
| **Study 2** | nono2T3 | No inhibition | 16 |
| | noN2T4 | Ammonia inhibition | 23 |
| | noN2T8 | Ammonia inhibition | 60 |
| | noN2T9 | Ammonia inhibition | 85 |
| | noPhi2T4 | Phenol inhibition | 23 |
| | noPhi2T5 | Phenol inhibition | 31 |
| | noPhi2T7 | Phenol inhibition | 50 |
| | AC1no2T3 | No inhibition | 16 |
| | AC1N2T4 | Ammonia inhibition | 23 |
| | AC1N2T6 | Ammonia inhibition | 39 |
| | AC1N2T9 | Ammonia inhibition | 85 |
| | AC1Phi2T3 | No inhibition | 16 |
| | AC1Phi2T5 | No inhibition | 31 |
| | AC2no2T3 | No inhibition | 16 |
| | AC2N2T11 | Ammonia inhibition | 119 |
| | AC2N2T4 | Ammonia inhibition | 23 |
| | AC2N2T8 | Ammonia inhibition | 60 |
| | AC2Phi2T3 | No inhibition | 16 |
| | AC2Phi2T5 | No inhibition | 31 |
| | Xno2T3 | No inhibition | 16 |
| | XN2T11 | Ammonia inhibition | 119 |
| | XN2T4 | Ammonia inhibition | 23 |
| | XN2T8 | Ammonia inhibition | 60 |
| | XPhi2T4 | Phenol inhibition | 23 |
| | XPhi2T5 | Phenol inhibition | 31 |
| | XPhi2T7 | Phenol inhibition | 50 |
| | Z1no2T3 | No inhibition | 16 |
| | Z1N2T4 | Ammonia inhibition | 23 |
| | Z1N2T6 | Ammonia inhibition | 39 |
| | Z1N2T8 | Ammonia inhibition | 60 |
| | Z1Phi2T4 | Phenol inhibition | 23 |
| | Z1Phi2T6 | Phenol inhibition | 39 |
| | Z2no2T3 | No inhibition | 16 |
| | Z2N2T4 | Ammonia inhibition | 23 |
| | Z2N2T6 | Ammonia inhibition | 39 |
| | Z2Phi2T4 | Phenol inhibition | 23 |
| | Z2Phi2T6 | Phenol inhibition | 39 |

| | | | |
|---|---|---|---|
| **Study 3** | SRR1507064_ | No inhibition | 15 |
| | SRR1507067_ | No inhibition | 15 |
| | SRR1507069_ | No inhibition | 15 |
| | SRR1507183_ | No inhibition | 15 |
| | SRR1507184_ | No inhibition | 15 |
| | SRR1507185_ | No inhibition | 15 |
| | SRR1507186_ | No inhibition | 15 |
| | SRR1507187_ | No inhibition | 15 |
| | SRR1507188_ | No inhibition | 15 |
| | SRR1507189_ | No inhibition | 15 |
| | SRR1507190_ | No inhibition | 15 |
| | SRR1507191_ | No inhibition | 15 |
| | SRR2296764_ | Ammonia moderate concentration | 15 |
| | SRR2296765_ | Ammonia moderate concentration | 15 |
| | SRR2296766_ | Ammonia moderate concentration | 15 |
| | SRR2296767_ | Ammonia moderate concentration | 15 |
| | SRR2296768_ | Ammonia inhibition, final days | 15 |
| | SRR2296769_ | Ammonia inhibition, early days | 15 |
| | SRR2296770_ | Ammonia inhibition, early days | 15 |
| | SRR2296771_ | Ammonia inhibition, early days | 15 |
| | SRR2296772_ | Ammonia inhibition, early days | 15 |
| | SRR2296773_ | Ammonia inhibition, final days | 15 |
| | SRR2296774_ | Ammonia inhibition, final days | 15 |
| | SRR2296775_ | Ammonia inhibition, final days | 15 |
| | SRR2296776_ | Ammonia inhibition, early days | 15 |
| | SRR2296777_ | Ammonia inhibition, early days | 15 |
| | SRR2296778_ | Ammonia inhibition, early days | 15 |
| | SRR2296779_ | Ammonia inhibition, final days | 15 |
| | SRR2296780_ | Ammonia inhibition, final days | 15 |
| | SRR2296781_ | Ammonia inhibition, final days | 15 |
| | SRR2296782_ | Ammonia inhibition, early days | 15 |
| | SRR2296783_ | Ammonia inhibition, early days | 15 |
| | SRR2296784_ | Ammonia inhibition, early days | 15 |
| | SRR2296785_ | Ammonia inhibition, final days | 15 |
| | SRR2296786_ | Ammonia inhibition, final days | 15 |
| | SRR2296787_ | Ammonia inhibition, final days | 15 |
| **Study 4** | SRR3629052 | No inhibition | 39 |
| | SRR3629054 | No inhibition | 99 |
| | SRR3629058 | No inhibition | 127 |
| | SRR3629059 | Ammonia inhibition start | 139 |
| | SRR3629149 | Ammonia inhibition start | 152 |
| | SRR3629150 | Ammonia inhibition | 172 |
| | SRR3629151 | Ammonia inhibition | 189 |
| | SRR3629152 | Ammonia inhibition | 212 |
| | SRR3629153 | Ammonia inhibition decrease | 223 |
| | SRR3629154 | Ammonia inhibition decrease | 232 |

**Table S2: taxonomic affiliation of the OTUs selected by MINT analysis (three inhibitors)**

| OTU name | Domain | Phylum | Class | Order | Family | Genus | Species |
|---|---|---|---|---|---|---|---|
| OTU_13 | Bacteria | Bacteroidetes | Bacteroidia | Bacteroidales | Bacteroidaceae | Bacteroides | Multi-affiliation |
| OTU_49 | Bacteria | Firmicutes | Clostridia | Clostridiales | Lachnospiraceae | Mobilitalea | unknown species |
| OTU_16 | Bacteria | Firmicutes | Clostridia | Clostridiales | Syntrophomonadaceae | Syntrophomonas | unknown species |
| OTU_29 | Bacteria | Bacteroidetes | Bacteroidia | Bacteroidales | Porphyromonadaceae | Proteiniphilum | unknown species |
| OTU_45 | Bacteria | Firmicutes | Clostridia | Clostridiales | Syntrophomonadaceae | Syntrophomonas | unknown species |
| OTU_31 | Bacteria | Firmicutes | Clostridia | Clostridiales | Lachnospiraceae | unknown genus | unknown species |
| OTU_75 | Bacteria | Firmicutes | Clostridia | Clostridiales | Family XI | Sporanaerobacter | Multi-affiliation |
| OTU_53 | Bacteria | Firmicutes | Clostridia | Clostridiales | Clostridiales vadinBB60 group | unknown genus | gut metagenome |
| OTU_71 | Bacteria | Firmicutes | Clostridia | Clostridiales | Ruminococcaceae | Multi-affiliation | Multi-affiliation |
| OTU_21 | Bacteria | Bacteroidetes | Bacteroidia | Bacteroidales | Porphyromonadaceae | Petrimonas | unknown species |
| OTU_26 | Bacteria | Chloroflexi | Anaerolineae | Anaerolineales | Anaerolineaceae | unknown genus | Multi-affiliation |
| OTU_11 | Bacteria | Cloacimonetes | W5 | unknown order | unknown family | unknown genus | unknown species |
| OTU_58 | Bacteria | Thermotogae | Thermotogae | Petrotogales | Petrotogaceae | AUTHM297 | unknown species |
| OTU_6 | Bacteria | Firmicutes | Clostridia | Clostridiales | Clostridiaceae 1 | Clostridium sensu stricto 1 | Multi-affiliation |
| OTU_4 | Archaea | Euryarchaeota | Methanomicrobia | Methanosarcinales | Methanosarcinaceae | Methanosarcina | Multi-affiliation |
| OTU_35 | Bacteria | Cloacimonetes | W5 | unknown order | unknown family | unknown genus | unknown species |
| OTU_46 | Bacteria | Firmicutes | Clostridia | Clostridiales | Syntrophomonadaceae | Syntrophomonas | Multi-affiliation |
| OTU_77 | Bacteria | Cloacimonetes | Cloacimonetes Incertae Sedis | unknown order | unknown family | Candidatus Cloacamonas | unknown species |
| OTU_102 | Bacteria | Firmicutes | Clostridia | Clostridiales | Syntrophomonadaceae | Syntrophomonas | unknown species |
| OTU_22 | Bacteria | Firmicutes | Clostridia | Clostridiales | Syntrophomonadaceae | Syntrophomonas | unknown species |
| OTU_156 | Bacteria | Firmicutes | Clostridia | Clostridiales | Clostridiaceae 1 | Caloramator | unknown species |
| OTU_14 | Bacteria | Bacteroidetes | Bacteroidia | Bacteroidales | Marinilabiaceae | Ruminofilibacter | Multi-affiliation |
| OTU_54 | Bacteria | Spirochaetae | Spirochaetes | Spirochaetales | Spirochaetaceae | Treponema 2 | unknown species |
| OTU_109 | Bacteria | Firmicutes | Bacilli | TSCOR001-H18 | unknown family | unknown genus | unknown species |
| OTU_96 | Bacteria | Bacteroidetes | Bacteroidia | Bacteroidales | Bacteroidaceae | Bacteroides | unknown species |
| OTU_86 | Bacteria | Firmicutes | Clostridia | Clostridiales | Peptostreptococcaceae | Peptostreptococcus | Multi-affiliation |
| OTU_27 | Bacteria | Firmicutes | Clostridia | Clostridiales | Family XI | Anaerosalibacter | unknown species |
| OTU_125 | Bacteria | Bacteroidetes | Bacteroidia | Bacteroidales | Bacteroidaceae | Bacteroides | unknown species |
| OTU_114 | Bacteria | Firmicutes | Clostridia | Clostridiales | Lachnospiraceae | Mobilitalea | unknown species |
| OTU_205 | Bacteria | Firmicutes | Clostridia | Clostridiales | Lachnospiraceae | Lachnospiraceae NK4A136 group | unknown species |
| OTU_139 | Bacteria | Firmicutes | Clostridia | Clostridiales | Caldicoprobacteraceae | Caldicoprobacter | Multi-affiliation |
| OTU_10 | Bacteria | Spirochaetae | Spirochaetes | Spirochaetales | Spirochaetaceae | Treponema 2 | unknown species |
| OTU_5 | Bacteria | Bacteroidetes | Bacteroidia | Bacteroidales | Porphyromonadaceae | Petrimonas | Multi-affiliation |
| OTU_33 | Bacteria | Firmicutes | Clostridia | Clostridiales | Defluviitaleaceae | Defluviitalea | unknown species |
| OTU_142 | Bacteria | Firmicutes | Clostridia | Clostridiales | Caldicoprobacteraceae | Caldicoprobacter | unknown species |
| OTU_202 | Bacteria | Firmicutes | Clostridia | Clostridiales | Family XI | Anaerosalibacter | Multi-affiliation |
| OTU_293 | Bacteria | Firmicutes | Clostridia | Clostridiales | Lachnospiraceae | unknown genus | unknown species |
| OTU_159 | Bacteria | Firmicutes | Bacilli | Lactobacillales | Enterococcaceae | Vagococcus | Multi-affiliation |
| OTU_74 | Bacteria | Bacteroidetes | Bacteroidia | Bacteroidales | Bacteroidaceae | Bacteroides | Multi-affiliation |
| OTU_115 | Bacteria | Firmicutes | Clostridia | Clostridiales | Family XI | unknown genus | unknown species |
| OTU_88 | Bacteria | Firmicutes | Clostridia | MBA03 | unknown family | unknown genus | unknown species |
| OTU_91 | Bacteria | Firmicutes | Clostridia | Clostridiales | Family XI | Tissierella | unknown species |
| OTU_155 | Bacteria | Firmicutes | Clostridia | Clostridiales | Clostridiaceae 1 | Clostridium sensu stricto 15 | Multi-affiliation |
| OTU_173 | Bacteria | Firmicutes | Clostridia | Clostridiales | Family XI | Tepidimicrobium | Clostridiales bacterium mt11 |
| OTU_12 | Bacteria | Bacteroidetes | Bacteroidia | Bacteroidales | Porphyromonadaceae | Proteiniphilum | unknown species |

**Table S3: taxonomic affiliation of the clusters (OTUs aggregated at the genus level) selected by MINT analysis (two inhibitors)**

| Group name | Domain | Phylum | Class | Order | Family | Genus |
|---|---|---|---|---|---|---|
| Cluster_15 | Bacteria | Firmicutes | Clostridia | Clostridiales | Defluviitaleaceae | Defluviitalea |
| Cluster_146 | Bacteria | Firmicutes | Clostridia | Clostridiales | Family XI | Tepidimicrobium |
| Cluster_87 | Bacteria | Firmicutes | Clostridia | Clostridiales | Family XI | unknown genus |
| Cluster_85 | Bacteria | Firmicutes | Clostridia | Clostridiales | Family XI | Tissierella |
| Cluster_110 | Bacteria | Firmicutes | Clostridia | Clostridiales | Caldicoprobacteraceae | Caldicoprobacter |
| Cluster_25 | Bacteria | Firmicutes | Clostridia | Clostridiales | Family XI | Anaerosalibacter |
| Cluster_132 | Bacteria | Spirochaetae | Spirochaetes | Spirochaetales | Spirochaetaceae | Sphaerochaeta |
| Cluster_69 | Bacteria | Firmicutes | Clostridia | Clostridiales | Ruminococcaceae | Multi-affiliation |
| Cluster_56 | Bacteria | Bacteroidetes | Bacteroidia | Bacteroidales | Porphyromonadaceae | Multi-affiliation |
| Cluster_43 | Bacteria | Cloacimonetes | Multi-affiliation | Multi-affiliation | Multi-affiliation | Multi-affiliation |
| Cluster_6 | Bacteria | Firmicutes | Clostridia | Clostridiales | Clostridiaceae 1 | Clostridium sensu stricto 1 |
| Cluster_34 | Bacteria | Firmicutes | Clostridia | Clostridiales | Ruminococcaceae | Ruminococcus 1 |
| Cluster_59 | Bacteria | Cloacimonetes | Cloacimonetes Incertae Sedis | unknown order | unknown family | Candidatus Cloacamonas |
| Cluster_42 | Bacteria | Firmicutes | Clostridia | Clostridiales | Christensenellaceae | Christensenellaceae R-7 group |
| Cluster_134 | Bacteria | Bacteroidetes | Bacteroidia | Bacteroidales | Porphyromonadaceae | unknown genus |
| Cluster_11 | Bacteria | Cloacimonetes | W5 | unknown order | unknown family | unknown genus |
| Cluster_17 | Bacteria | Firmicutes | Clostridia | Clostridiales | Syntrophomonadaceae | Syntrophomonas |
